# Embryonically generated interneurons (EGINs) exhibit a distinctive developmental program and lamination in the olfactory bulb

**DOI:** 10.1101/2024.11.25.625264

**Authors:** Natalie Spence, Eduardo Martin-Lopez, Kimberly Han, Marion Lefèvre, Nathaniel W. Lange, Bowen Brennan, Charles A. Greer

## Abstract

Interneurons (INs) in the mouse olfactory bulb (OB) develop during an extensive period of time that begins in the embryo and continues throughout the lifetime of an individual. The connectivity and location of these neurons is profoundly affected by the time of generation, making these developmental windows critical to understand the circuitry of the OB. Here, we focus on the OB embryonic generated interneurons (or EGINs), which have generally received less attention than those generated in the adult. Birthdates of EGINs were differentiated by embryonic injections of thymidine analogs and their final destinations and phenotypes analyzed by immunohistochemistry. We found that EGINs were retained in the adult and were distributed across all layers of the OB. However, a lateral-to-medial neurogenic gradient is seen only in the earliest generated EGINs. Within the granule cell layer (GCL), EGINs predominantly accumulated in the superficial region at almost all ages and remained there during adulthood. Immunostaining for calbindin, parvalbumin, tyrosine hydroxylase, and calretinin were largely negative suggesting that EGINs may represent subpopulations of OB INs that are not yet fully characterized. Using in utero electroporation to label EGIN progenitors, we demonstrated that they reach the primordial OB as early as E13 and begin to differentiate apical dendrites by E15. At E16, EGIN progenitors clustered into a chain of migrating neuroblasts denoting the embryonic rostral migratory stream (RMS). Collectively, our data highlight the importance of studying OB INs in isolated time windows to better understand the formation of circuits that define the olfactory responses.

## Introduction

The olfactory bulb (OB), together with the hippocampus, is one of the regions of the brain that retains adult neurogenesis receiving a constant influx of neuroblasts during the lifetime of an individual (Bayer, 1985; Whitman and Greer, 2009). Neuroblasts targeted to the OB are generated in the subventricular zone (SVZ) and migrate via the rostral migratory stream (RMS) to reach the OB, where they differentiate into several classes of interneurons (INs) (Lois and Alvarez-Buylla, 1994; Lledo et al., 2008). Although this is the pathway followed by OB neuroblasts from birth, the production of OB INs begins embryonically in the dorsal part of the lateral ganglionic eminence (LGE). The vast majority of OB INs are generated at young postnatal ages and decline to only about 2% during adulthood (Kim et al., 2020). The continuous addition of INs into the OB circuitry turns the OB into one of the most plastic environments of the brain, which is crucial for essential behaviors such as feeding, social interactions, reproduction, memory formation, and many other behaviors (Lazarini and Lledo, 2011; Sakamoto et al., 2011; Naffaa, 2025). This process may reflect an evolutionary response to unexpected and diverse stimuli throughout development and life (Alvarez-Buylla et al., 2001). Disruptions in early IN development is seen in various neurodevelopmental disorders, including autism and Tourette’s syndrome, and has implications for mental health (Lyons-Warren et al., 2021; Meller et al., 2023; Dejou et al., 2024).

The OB has a very high ratio of INs to projection neurons (100:1) compared to any other brain structure (Shepherd et al., 2021). OB INs are small, often anaxonic, and modulate the activity of the OB projection neurons (Whitman and Greer, 2007a). Depending on their soma location across the OB layers, the OB INs can be classified in four groups (Nagayama et al., 2014). The first are the periglomerular cells (PGCs), located in the vicinity of the glomerular layer (GL) and make connections with olfactory sensory neuron axons from the epithelium. They also make dendrodendritic connections with the OB projection neurons, mitral and tufted cells (M/Tc) (Wachowiak and Shipley, 2006). Second, INs of the external plexiform layer (EPL) make reciprocal connections with the mitral cells, which include the superficial short-axon neurons (sSA) and multipolar neurons, among others (Nagayama et al., 2014). The third are superficial granule cells (GCs) located in the mitral cell layer (MCL), which account for the majority of cells in this layer (Greer et al., 2008). Fourth, are GCs in the granule cell layer (GCL), which contains the vast majority of OB INs at ∼94%, and project their dendrites into the EPL (Greer et al., 2008; Nagayama et al., 2014). It is generally agreed that these distinct subpopulations of INs contribute differentially to synaptic connectivity and activity in the OB (Mori and Sakano, 2021).

An important aspect of the OB INs is that their function and integration into circuits is impacted by the time of generation, particularly between embryonic and postnatal ages (Fritz et al., 1996; Batista-Brito et al., 2008; Galliano et al., 2018; Takahashi et al., 2018; Kim et al., 2020). This makes the neuronal birthdate a fundamental factor in regulating the organization and development of the OB. Similarly, INs generated within the same developmental periods postnatally have distinct properties and phenotypes. INs generated early in postnatal development compared with those generated at later times differ in their circuitry and lamination (Lemasson et al., 2005; Muthusamy et al., 2017). Most recently, this has been explored by our lab where timing of neurogenesis in perinatal mice influences the placement of GCs in the OB (Liberia et al., 2024). Similarly, the projection neurons in the OB (M/Tc), are distinguished based on the timeframe of neurogenesis with early and late born cells differentially distributed in the OB and with stereotyped projections to olfactory cortex, compared to earlier generated ones that extend throughout the whole cortex (Imamura et al., 2011; Imamura and Greer, 2015; Chon et al., 2020).

While adult neurogenesis in the OB is widely acknowledged, we sought here to focus on the development of EGINs in the OB in order to assess that the timing of generation influences their placement in the OB across embryonic development and into adulthood. We tracked neurogenesis using thymidine analogs and immunohistochemistry. To assess EIGN migration we performed in utero injection and electroporation (IUE), as we previously used in the olfactory cortex, tubular striatum, and perinatal OB (Martin-Lopez et al., 2019a; Martin-Lopez et al., 2019b; Liberia et al., 2024). We established the final position of EGINs in the adult OB following migration and revealed an initial lateral-to-medial developmental gradient. Furthermore, our data support that EGINs in the GCL generated at the earliest embryonic ages tended to occupy the outer region of the GCL prior to the deeper regions. These data led to the conclusion that the connectivity of EIGNs differs among OB projection neurons based on time of origin. We also analyzed EGIN phenotypes at their final destinations in the adult OB, finding that they did not belong to those most known subpopulations of INs, identified by common antibodies (Batista-Brito et al., 2008). We compared these phenotypes with INs generated in juvenile postnatal mice (PGINs), which gave rise to similar results. Finally, we studied the morphology and migration of EGINs using IUE, which allowed us to examine the migratory pathways, dendritogenesis, and RMS formation at embryonic stages. Collectively, this analysis shows the importance of timing of embryonic neurogenesis in relation to anatomical position and cell characterization in the adult OB.

## Materials and Methods

### Animals

All experiments were performed using pregnant and postnatal CD1 mice (Charles River, cd-1r-igs) at different embryonic stages. In analyzing embryonic timepoints, embryonic day 0 (E0) was considered as the day that the vaginal plug was found. Mice were maintained on a 12h light cycle and received chow and water *ad libtium* in the vivarium at Yale University. All protocols and procedures were approved by Yale University Animal Care and Use Committee.

### Thymidine analogs injection

To assess the date of embryonic generation of EGINs, two different thymidine analogs were injected per dam to differentiate between two embryonic ages in the same animals as previously described (Martin-Lopez et al., 2019a). Pregnant females received double intraperitoneal (IP) injections (separated by 2 hours) of 50 mg/kg of 5-Iodo-2′-deoxyuridine (IdU; Sigma Aldrich, I7125) followed by 50 mg/kg 5-Chloro-2′-deoxyuridine (CldU; Sigma-Aldrich, C6891) four days apart, using the following sequence: E10-E14 / E11-E15 / E12-E16 / E13-E17 / and only E18 (IdU). EGIN lamination in the OB were studied at postnatal day 21 (P21). To compare the phenotype of these cells with those generated postnatally in juvenile animals, a second set of mice were injected at postnatal day 25 (P25) with 50 mg/kg of 5-Bromo-2’-Deoxyuridine (BrdU; RPI Research Products, B74200-0.5) diluted in Dulbecco’s Phosphate Buffered Saline (DPBS, Gibco 14190-144). These cells, that we referred to as postnatally generated interneurons (PGINs), were studied 25 days after BrdU injection at P50.

### In utero injection and electroporation (IUE)

EGINs are reported to arise within the lateral ganglionic eminence (LGE) (Toresson and Campbell, 2001; Wichterle et al., 2001; Guo et al., 2019). To study the migration and differentiation of EGINs, we labeled their progenitor cells using a multicolor piggyBac technique that target the LGE using IUE (Martin-Lopez et al., 2019a).,A mixture of four plasmids was prepared that contained three different piggyBac transposons expressing the fluorophores: pPB-CAG-EGFP (for green), pPB-CAG-tdTomato (for red), and pPB-CAG-iRFP670 (for far red, but pseudo colored in blue), combined with a plasmid expressing the enzyme transposase (pCAG-PBase). All plasmids were prepared at a final concentration of 1 μg/μL supplemented with 0.05% of fast green (Sigma-Aldrich, F7252) for visualization during the injections. This mixture was loaded into borosilicate capillaries prior to injection. The procedure began by anesthetizing pregnant females at the E11 embryonic stage with 2.5% isoflurane, placed at the supine position over a heating pad, and the belly shaved and disinfected in preparation for the procedure. An incision was made along the midline in the abdominal wall, and the peritoneal membrane was cut through the alba line. Embryos were extracted from the abdominal cavity and injected in the lateral ventricles using the glass capillaries filled with the plasmid mixture connected to a Picospritzer (General Valve Corporation). Immediately after, the electroporation of the LGE was achieved by delivering 5 pulses of 35 V using a pair of gold tweezers (Genepaddles-542, Harvard Apparatus, 45-0122) connected to an ECM 830 electroporator (BTX Harvard Apparatus). After the embryos were placed back inside the abdominal cavity, the peritoneal membrane was closed using 5/0 PGA absorbable sutures (AD Surgical, S-G518R13-U), and the skin sutured using 5/0 silk braided sutures (AD Surgical, S-S518R13). Postsurgical care involved the administration of 4 mg/kg of the analgesic Meloxicam (Covetrus, 049756) injected subcutaneously for 72h.

### Tissue processing and immunostaining

Mice that received injections of thymidine analogs were studied at the ages of P21 for EGINs and P50 for PGINs, respectively. Mice were euthanized using an overdose of Euthasol (Virbac) and then transcardially perfused with 5 mL of saline solution, followed by 20 mL of 4% paraformaldehyde (PFA) in phosphate-buffered saline (PBS) at 4°C. Afterwards, brains were dissected from the skulls and post-fixed in PFA overnight. Two groups of IUE mice were used: those whose brains were analyzed postnatally were processed identically as described above; and those studied embryonically, whose tissues were extracted from CO_2_ euthanized damns at different ages. These embryonic brains were fixed by immersion in PFA for 2 days at 4°C. Finally, all brains were cryoprotected by immersion in a 30% sucrose solution and embedded in Tissue-Tek OCT compound (Fisher Scientific, catalog #4585) to prepare blocks for cryosectioning. Brains were serially sectioned at 25µm in the coronal plane using a Reichert Frigocut cryostat (E-2800), then dried out using a slide warmer at 60°C and stored at -80 °C.

In preparation for immunochemistry, slides were thawed on a slide warmer at 60°C for 30 minutes. To detect thymidine analogs, nuclear DNA was denatured by treating the sections for 35 minutes in 0.025 M HCl at 65°C, followed by a 5-minute heat-shock in ice-cooled 0.025 M HCL solution. The HCL was rinsed and neutralized with 0.1 M borate buffer (pH 8.5) and washed with PBS supplemented with 0.1% Triton X100 (American Bioanalytical, AB02025) (PBST). Slides were then blocked in PBST supplemented with 5% Normal Goat Serum (NGS; Accurate Chemicals, CL1200-100) and 0.1% Bovine Serum Albumin (BSA; Sigma-Aldrich, A3059) for one hour at room temperature (RT). After this, primary antibodies (Table 1) were diluted in a 1:10 dilution of blocking solution, added to the sections, and then incubated overnight at 4°C. The following morning the slides were washed 3 times for 10 minutes each with PBST, and then incubated with the appropriate secondary antibodies (Table 1) diluted in PBST and supplemented with DAPI (Sigma-Aldrich, D9542) for nuclear counterstaining, for 2 hours at RT. To ensure that quantified cells were not astrocytes or microglia, additional staining was performed against glial markers (Table 1). Finally, NeuN staining was done to confirm the IdU/CldU/BrdU cells were neurons (Fig. 1).

**Table 1.** Antibodies.

| Antigen | Primary Ab | Source (Cat. #) | RRID | Dilution | Secondary Ab | Source | Dilution |
| --- | --- | --- | --- | --- | --- | --- | --- |
| BrdU/IdU | Mouse IgG1 | BD Biosciences (347580) | AB_400326 | 1:200 | Goat α-mouse IgG (H+L)-Alexa 555 Supercloonal™ Recombinant | Thermo Fisher Scientific | 1:1000 |
| BrdU/CldU | Rat IgG2a | Abcam (ab6326) | AB_305426 | 1:300 | Goat α-rat IgG Alexa 488 | Thermo Fisher Scientific | 1:1000 |
| Calbindin D28K | Rabbit | Millipore (MAB1778) | AB_2068336 | 1:500 | Goat α-rabbit IgG Alexa 647 | Thermo Fisher Scientific | 1:1000 |
| Calretinin | Mouse IgG1 | Millipore (MAB1568) | AB_94259 | 1:1000 | Goat α-mouse IgG Alexa 647 | Thermo Fisher Scientific | 1:1000 |
| GFAP | Rabbit IgG | Biologend (840001) | AB_2565444 | 1:500 | Goat α-rabbit IgG Alexa 647 | Thermo Fisher Scientific | 1:1000 |
| IBA1 | Rabbit IgG | Wako (016-200001) | AB_839506 | 1:500 | Goat α-rabbit IgG Alexa 647 | Thermo Fisher Scientific | 1:1000 |
| NeuN | Rabbit IgG | GeneTex (GTX132974) | AB_2886793 | 1:500 | Goat α-rabbit IgG Alexa 647 | Thermo Fisher Scientific | 1:1000 |
| Parvalbumin | Rabbit IgG | Abcam (ab11427) | AB_298032 | 1:100 | Goat α-rabbit IgG Alexa 647 | Thermo Fisher Scientific | 1:1000 |
| Reelin (recombinant) | Rabbit IgG | Abcam (ab312310) | AB_3076463 | 1:100 | Goat α-rabbit IgG Alexa 647 | Thermo Fisher Scientific | 1:1000 |
| Tyrosine Hydroxylase | Mouse IgG1 | Millipore (MAB318) | AB_2201528 | 1:400 | Goat α-mouse IgG Alexa 647 | Thermo Fisher Scientific | 1:1000 |

### Imaging, quantifications, and statistical analysis

Images for IdU/CldU quantification were acquired using a 10X objective in an Olympus BX51 epifluorescence microscope. Images for BrdU quantification were obtained using a laser scanning confocal microscope (Zeiss LSM 900 with Airyscan) using a 20X objective. Images from IUE mice were taken with a laser scanning confocal microscope (Zeiss LSM 800 with Airyscan) in the developing OB.

**Figure 1.**
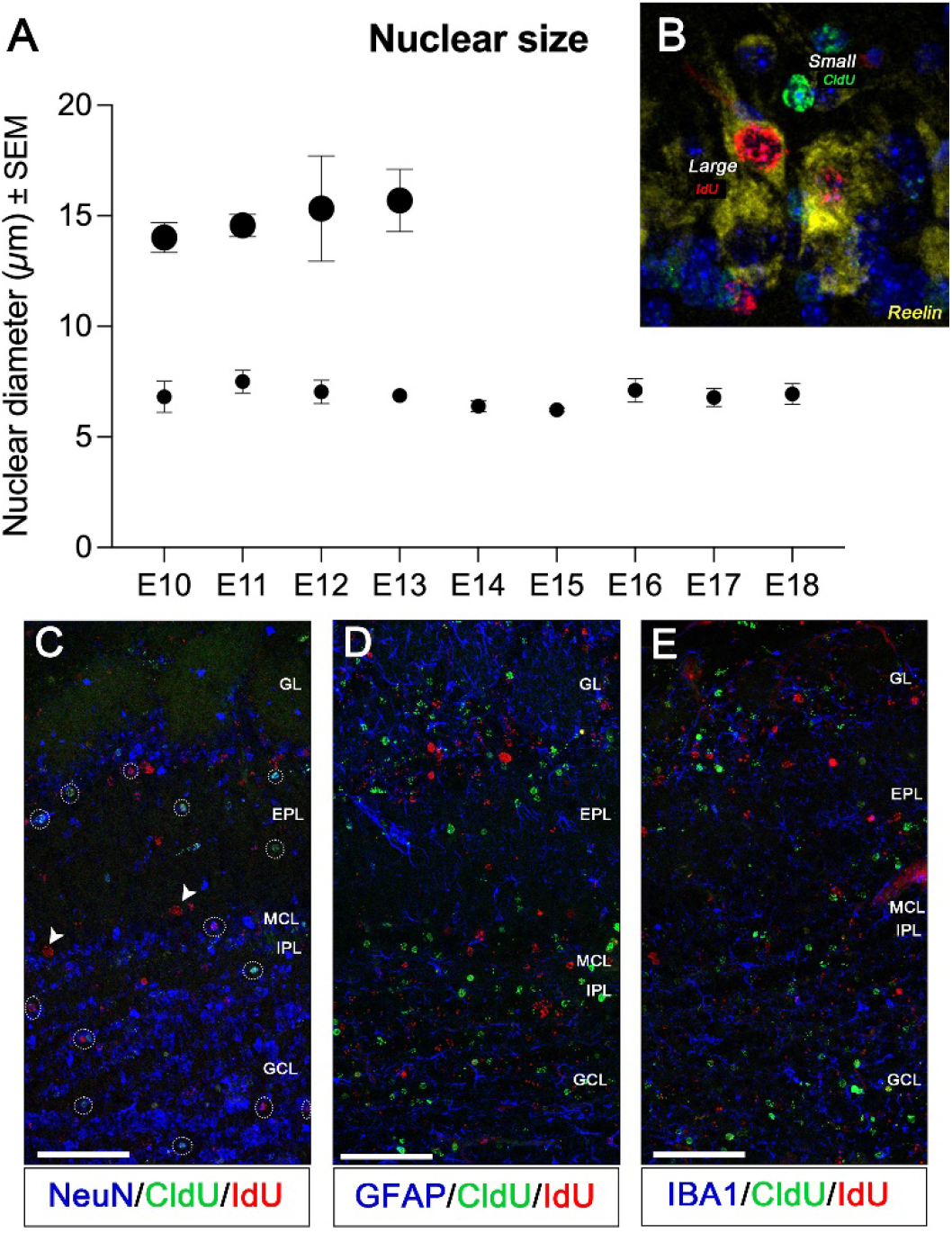
EGINs identification in the OB. **(A)** EGINs identification by nuclear size. Representation of the average diameters from nuclear size between “small” nuclei (EGINs) and “large” nuclei (projection neurons). (**B)** Depicts size difference between the nuclei sizes and colocalization with reelin (yellow), a marker for OB projection neurons. (**C)** Immunostaining for NeuN, a marker for mature interneurons in the OB, is shown colocalizing (dashed circles) with the thymidine analogs CldU and IdU. Projection neurons are not expressing NeuN, indicated by arrows. (**D** and **E)** Immunostaining showing a lack of colocalization with GFAP and IBA1, which are markers for astrocytes and microglia, respectively. Abbreviations: CldU: 5-Chloro-2′-deoxyuridine; EPL: external plexiform layer; GCL: granule cell layer; GFAP: glial fibrillary acidic protein; GL: glomerular layer; IBA1: Ionized calcium-binding adaptor molecule; IdU: 5-Iodo-2′-deoxyuridine; MCL/IPL: mitral cell layer/internal plexiform layer; NeuN: neuron specific nuclear protein. Scale bars: 100 μm.

To avoid quantifying projection neurons, which also develop at early embryonic stages (Blanchart et al., 2006; Imamura et al., 2011), we analyzed individual nuclei and measured their diameters since it is known that OB projection neurons are larger in size (>15 μm diameter) than INs (5-15 μm diameter) (Nagayama et al., 2014). Nuclear diameters were measured using the ImageJ measuring tool and only that had a nuclei size that was under 10 µm in length (n=3/group) were considered INs. Further exclusion criteria for projection neurons was made by staining the section with the marker Reelin, known to specifically label M/Tc in the OB (Martin-Lopez et al., 2011).

For quantification of IdU/CldU positive cells (EGINs), anatomically matched images of three OB sections along the anterior-to-posterior axis were used. Among these sections, lateral and medial regions were defined. CldU^+^ and IdU^+^ cells were counted in four layers of the OB, including the glomerular layer (GL), external plexiform layer (EPL), mitral cell layer and internal plexiform layer (MCL/IPL) that were combined due to their proximity, and granule cell layer (GCL). Additionally, the GCL was divided into four subregions of 50 μm thickness each to quantify cells going from the superficial (Layer 1) to the deep (Layer 4) GCL. Using the regions of interest (ROIs) functionality in ImageJ, OB layer delineations were copied from image to image, allowing OB layer areas to be maintained across all quantification images. The total number of labeled cells generated at different embryonic timepoints were then counted along the medial-to-lateral axis and amongst OB layers using ImageJ Cell Counter in six sections per animal for each timepoint (*n =* 6 / 3 males and 3 females per group). The final numbers were then normalized as cells/mm^2^. A two-way ANOVA was used to compare the numbers of labeled cells between sexes across the lateral-medial axis and age. Since no significant differences were observed, male and female data were combined for further analysis. To analyze distribution of total labeled cells among OB layers along the lateral-to-medial axis and in relation to timing of neurogenesis, we performed unpaired t-tests and a two-way ANOVA with a post hoc Tukey test (GraphPad Prism 10.4.0).

To test for neuronal differentiation, we quantified the percentage of EGINs expressing one of the most commonly known markers distinctive of OB INs (Parrish-Aungst et al., 2007): calbindin (CB), parvalbumin (PV), tyrosine hydroxylase (TH) and calretinin (CR). Using the same ROIs as described above, we quantified the total number of cells expressing each marker by layer, and then the number of IdU^+^ and CldU^+^ cells that co-expressed each individual marker. These numbers were compared to those of postnatal mice to check neuronal differentiation influenced by generation before or after birth. Postnatal mice were injected with BrdU, and quantification of BrdU^+^ cells (PGINs) co-expressing the INs markers was done using a similar protocol as for EGINs. For PGINs, three serial sections from each OB along the anterior-posterior axis were used for the analysis, and BrdU^+^ cells were counted across the OB layers. Quantifications were done using ImageJ software and the numbers averaged by animal and normalized as cells/mm^2^. Colocalization data from EGINs and PGINs was represented as pie charts using GraphPad Prism 10.4.0.

## Results

### Characterization of EGINs nuclei in the adult olfactory bulb

OB INs are embryonically generated in the LGE followed by migration to the OB (Toresson and Campbell, 2001; Wichterle et al., 2001). These INs survive until adulthood, becoming a specialized subset of OB INs that we refer to as EGINs. Although EGINs were previously recognized (Hinds, 1968a; Alvarez-Buylla et al., 1994; Batista-Brito et al., 2008; Takahashi et al., 2018), the relationships between their generation dates with their location and molecular diversity in the adult OB is not yet fully understood. Here, we used thymidine analogs to analyze the EGINs birthdates and their placement in the adult OB.

In order to proceed with the analysis of EGIN birthdates and distribution, we first established a criterion of exclusion for projection neurons, since analogs of thymidine do not distinguish between cell types at the time of injection. EGINs and projection neurons share a generation window between E10-E13 (Hinds, 1968a; Blanchart et al., 2006; Batista-Brito et al., 2008). To distinguish EGIN nuclei from those of the projection neurons, we measured the diameter of the nuclei finding two clearly distinctive sets: those that measured an average of 7±0.4 μm in diameter that appeared at all studied ages (Fig. 1A); and those that measured an average of 15±0.6 μm in diameter that were only visible from E10-E13 (Fig. 1A). Because these latter diameters are consistent with prior descriptions of projection neurons (Nagayama et al., 2014), and fall into the appropriate generation window of M/Tc (Blanchart et al., 2006; Imamura et al., 2011), we decided to exclude nuclei larger than 10 μm from our analyses. Additionally, we performed immunohistochemistry against reelin, a marker typical of M/Tc. (Alcantara et al., 1998; Martin-Lopez et al., 2011), to exclude OB projection neurons. We found that only larger nuclei colocalized with reelin (Fig. 1B). To further confirm the neuronal nature of EGINs we co-stained with NeuN, a marker typically expressed by mature interneurons in the OB, to show that small-nuclei cells expressed NeuN (Fig 1C, circled nuclei). This staining was absent from the larger nuclei, as M/Tc do not express NeuN (Mullen et al., 1992) (Fig. 1C, arrowheads). The non-glial nature of these cells was confirmed by immunohistochemistry against the astrocyte marker glial fibrillary acid protein (GFAP), and the microglial marker ionized calcium binding adapter molecule 1 (IBA1), showing a complete lack of staining for these markers (Fig. 1D, E).

### EGINs across the lateral-to-medial axis of the olfactory bulb

We further analyzed the distribution of EGINs in the adult OB along the lateral-to-medial axis to assess for a developmental neurogenic gradient as previously demonstrated for mitral cells. Mitral cells generated earlier localized to the dorsomedial MCL, whereas late-generated cells localized to the ventrolateral MCL (Imamura et al., 2011). Here, to test for a lateral-to-medial neurogenic gradient of EGINs, we quantified nuclei in both sides of the OBs at postnatal day P21. As expected, we observed a robust cell labeling in all OB tissues receiving EGINs from all embryonic timepoints studied (Fig. 2A-E). The quantification across the lateral-to-medial axis revealed an initial preference towards the lateral side of the OB, starting at E11 and lasting until E13 (Fig. 2F and Table 2). From E14 onwards, this pattern dissipates and there was not a difference in the number of EGINs distributed on the lateral versus medial sides of the OB. Lateral-to-medial neurogenic patterns have been shown previously in other regions of the olfactory system such as the tubular striatum (a.k.a. the olfactory tubercle) (Martin-Lopez et al., 2019b). Additionally, prior to birth at E18, we observed a dramatic increase of EGINs in the OB that almost doubled the number of cells in the entire OB (Fig. 2F). This was seen across all layers (Fig. 3). This is consistent with previous reports demonstrating a spike in cells at the end of the gestation period in mouse (Hinds, 1968b), but in contrast with data from rats in which spikes in OB IN neurogenesis were not seen during embryonic stages (Bayer, 1983).

**Table 2.**
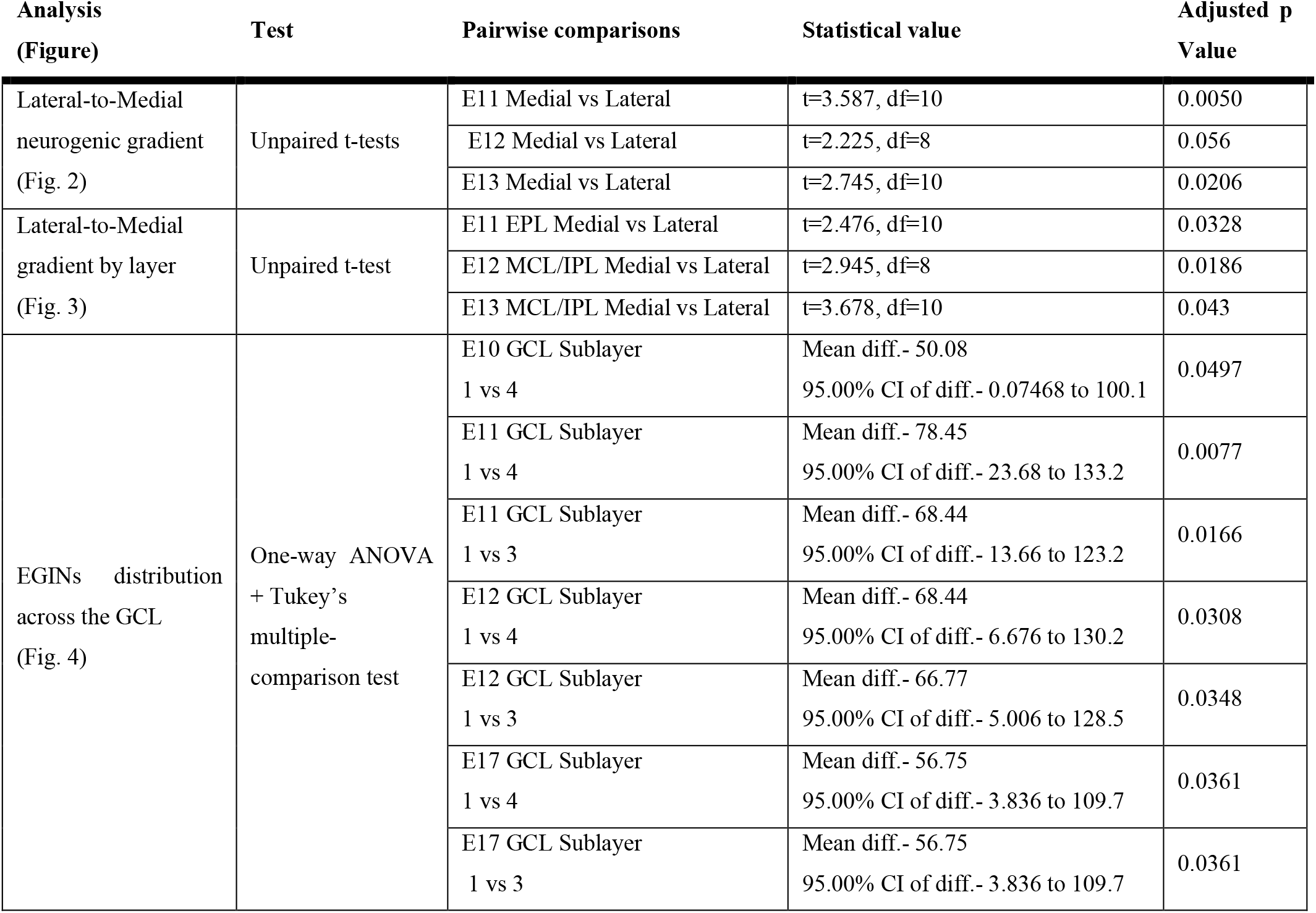
Statistical analysis.

**Figure 2.**
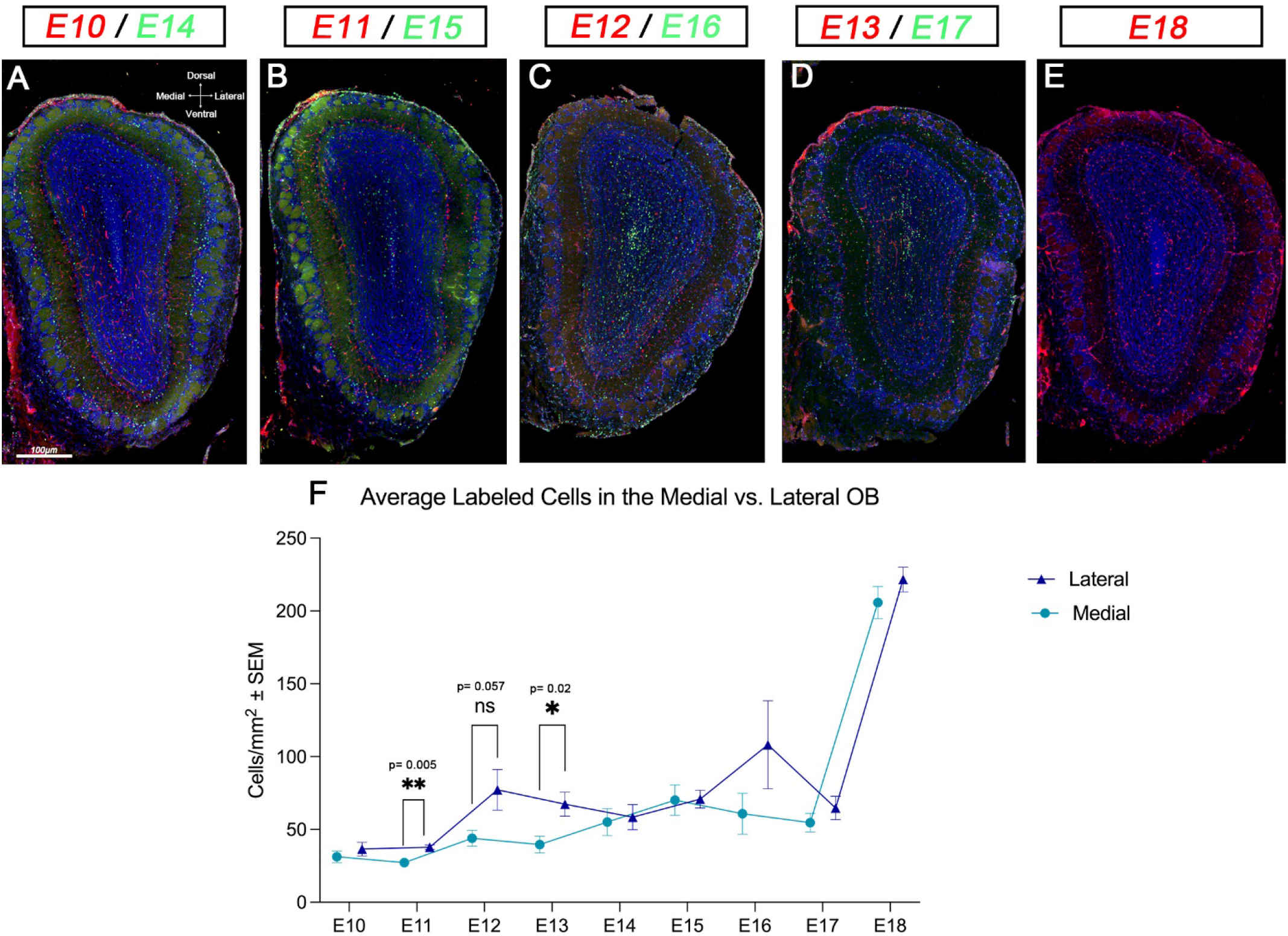
EGIN distribution across the OB. (**A-E**) Representative images of EGINs generated between the embryonic days E10 to E18 in the OB showing IdU+ cells (red) and CldU+ cells (green) with nuclei counterstained with DAPI (blue). (**F**) Quantification of EGINs in the OB across the lateral-to-medial axis from E10-E18. The graphic shows a significant increase in EGINs in the lateral sides of the OB at E11 and E13. An increase in the lateral side at E12 is shown but not statistically significant due to variability. On E18 there is a dramatic increase in EGINs in both the lateral and medial sides of the OB. Statistical significance: ^*^ = *p* < 0.05, ^**^ = *p* < 0.01. Scale bars: 100μm.

**Figure 3.**
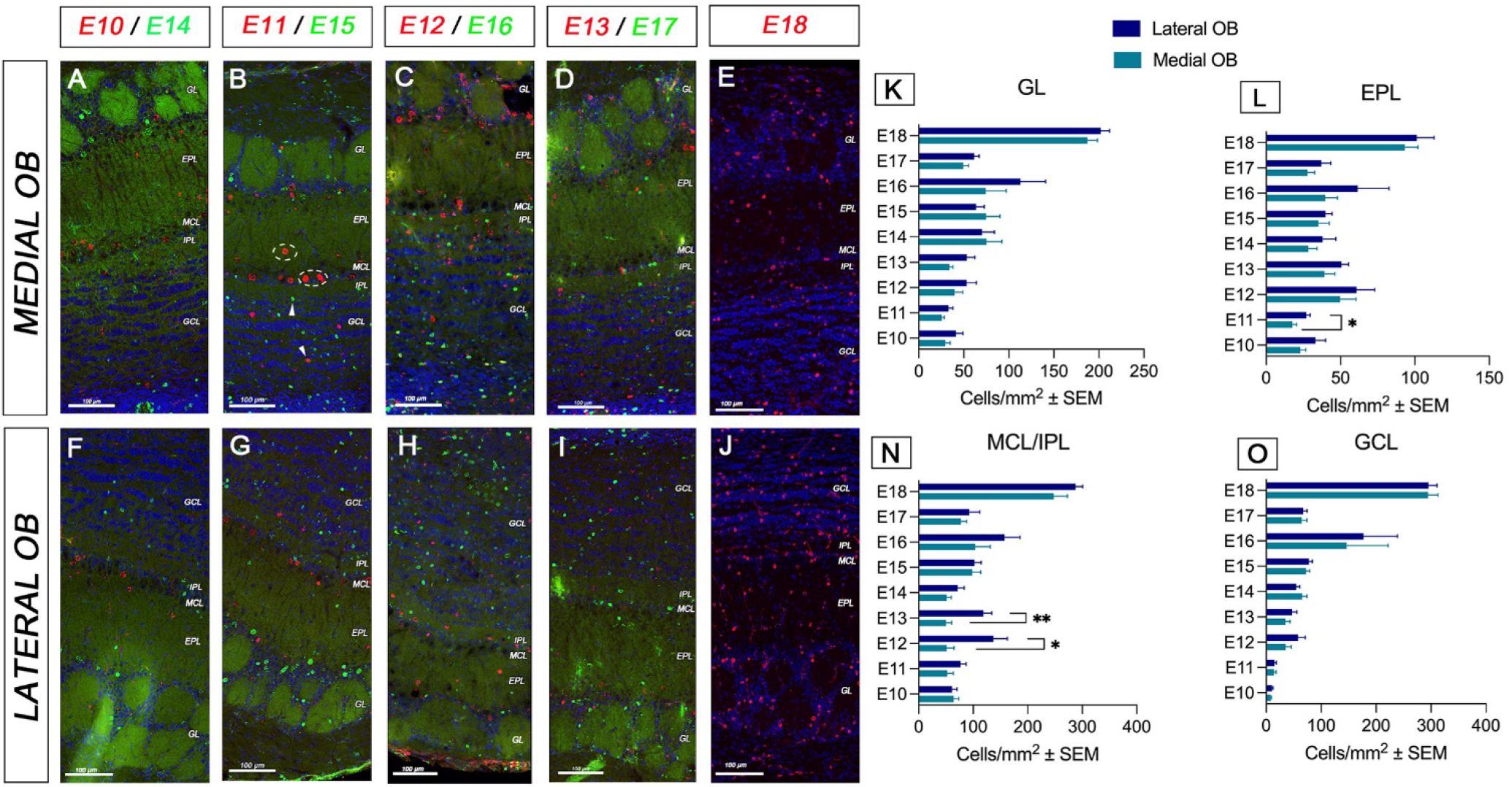
EGINs lateral-to-medial gradient across OB layers. Depict sections from **(A-E)** the medial OB and (**F-J)** from the lateral OB, showing IdU^+^ cells (red) and CldU^+^ cells (green), with nuclei counterstained with DAPI (blue). Arrows in **(B)** indicate interneurons, dashed circles indicate projection neurons. (**K-O)** Quantification of EGINs between the lateral and medial regions of the OB segregated by layers. Of interest are the statistical differences between lateral and medial distribution in the EPL at E11 (**L**), and at E12 and E13 in the MCL/IPL (**N**). Abbreviations: EPL: external plexiform layer; GCL: granule cell layer; GL: glomerular layer; MCL/IPL: mitral cell layer/internal plexiform layer. Statistical significance: ^*^ = *p* < 0.05, ^**^ = *p* < 0.01. Scale bars: 100 µm.

### Distribution of EGINs across OB layers

To understand if the E11-E13 lateral-to-medial gradient occurred across all or only some of the OB layers, we run our analysis by layer. Our data showed that that the lateral-to-medial gradient occurred in the EPL and MCL/IPL (Fig. 3). In the EPL, there was a significantly higher number of EGINs in the lateral region compared to the medial at E11 (Fig. 3L), indicating that the EPL medial axis to assess for a developmental neurogenic gradient as previously demonstrated for mitral cells. Mitral cells generated earlier localized to the dorsomedial MCL, whereas late-generated cells localized to the ventrolateral MCL (Imamura et al., 2011). Here, to test for a lateral-to-medial neurogenic gradient of EGINs, we quantified nuclei in both sides of the OBs at postnatal day P21. As expected, we observed a robust cell labeling in all OB tissues receiving largely accounted for the lateral-to-medial gradient throughout the OB (Fig. 2F). Similarly, we showed that the differences observed at E12 and E13 were caused by an increase in the lateral production of EGINs in the MCL/IPL (Fig. 3N), likely due to the superficial GCs (Greer et al., 2008).

To test if EGINs replicate the “superficial-first to deep-later” generation pattern seen in postnatal INs (Hinds, 1968b; Lemasson et al., 2005; Liberia et al., 2024), we divided the GCL into sublaminae. We quantified four 50 µm sublayers (Liberia et al., 2024), starting in the MCL where the superficial GCs are located (Layer 1 - superficial), and ending close to the RMS (Layer 4 - deep) (Fig. 4A). Our results showed that layer 1 consistently had the most EGINs across the ages E13-E17 (Fig. 4B-I), but then equaled to the other sublayers at E18 (Fig. 4J) where the influx of EGINs occurred (Fig. 2F). Although not always statistically significant, the pattern of early generated EGINs going towards the most superficial layer of the GCL (Layer 1) remained consistent throughout the entire embryonic development (Fig. 4B-I). The statistical significance highlighted that this pattern was more evident at early embryonic stages (Fig. 4B-D). These data supported the notion that the anatomical location of EGINs, as other GCs, was influenced by timing of neurogenesis in the OB (Hinds, 1968b; Batista-Brito et al., 2008; Galliano et al., 2018; Takahashi et al., 2018; Kim et al., 2020; Liberia et al., 2024).

**Figure 4.**
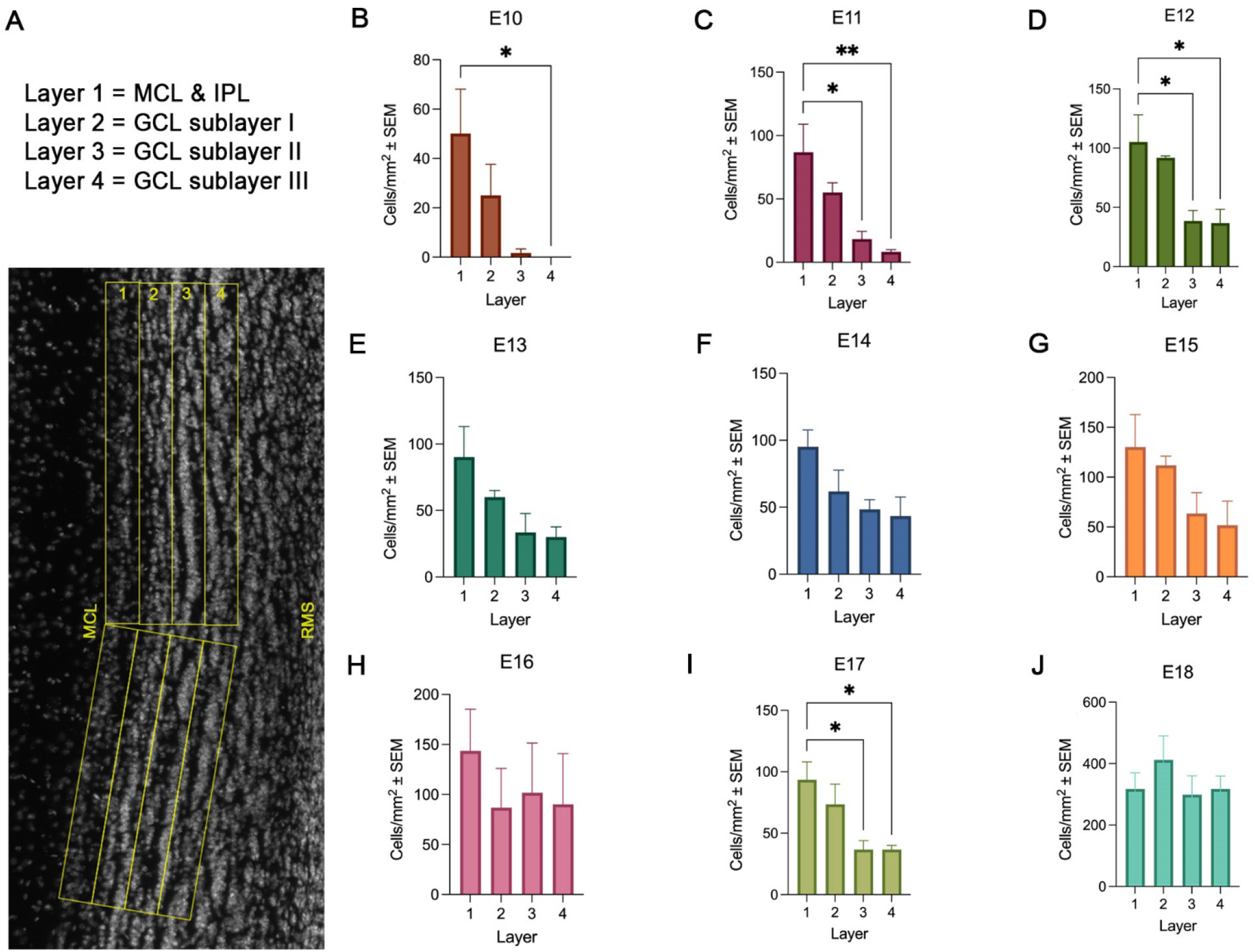
EGINs across GCL sublayers. (**A)** Template of the four sublayers within the GCL used for quantification over a DAPI stained OB section. Layer 1= MCL/IPL considered the most superficial and layer 4= deepest. Each sublayer is 50μm apart. (**B-J)** Quantification of EGINs showing a general preference for the superficial MCL/IPL layer 1 superficial layer versus the deep layer 4. In E10, E11, E12, and E17 there are significantly more EGINs in the superficial vs the deeper GCL layers. By E18, EGINs are more evenly distributed across the whole GCL. Abbreviations: GCL: granule cell layer; MCL/IPL: mitral cell layer/internal plexiform layer; RMS: rostral migratory stream. Statistical significance: ^*^ = *p* < 0.05, ^**^ = *p* < 0.01.

### Characterization of EGINs phenotypes using common interneuron markers

We next asked if EGINs grouped into well-defined immunohistochemically defined subpopulations of OB interneurons including: calbindin (CB), calretinin (CR), tyrosine-hydroxylase (TH), and parvalbumin (PV) (Parrish-Aungst et al., 2007; Batista-Brito et al., 2008; Nagayama et al., 2014). To obtain the percentage of EGINs expressing each marker, we first quantified each marker individually across the OB layers and characterized their location. As anticipated, we found the majority of CB (Fig. 5A-E, U) and TH cells (Fig. 5 K-O, W) located in the GL, PV in the EPL (Fig. 5F-J, V) and CR more widely distributed across the OB layers (Fig. 5P-T, X). To assess for EGIN differentiation, we quantified the number of cells that co-expressed these markers with the analogs of thymidine at P21. Interestingly, our data showed that only a small fraction of EGINs differentiated into the four populations of studied INs: CB: 0.3% (Fig. 5U), PV: 0.2% (Fig. 5V), TH: 1.4% (Fig. 5W) and CR: 2.8% (Fig. 5X). Based on this observation we hypothesized that INs expressing these four markers were generated less frequently at embryonic stages and instead were generated postnatally. We investigated this by injecting P25 mice with BrdU and analyzed the tissues 25 days later at P50. Surprisingly, our data showed that the percentage of these cell types generated postnatally remained low or inexistent: CB: 0% (Fig. 5U), 0: 0.2% (Fig. 5V), TH: 1.7% (Fig. 5W) and CR: 8.1% (Fig. 5X). These data suggested that the production and maturation of these cells occurred slowly over the course of postnatal development and likely takes longer than a month to maturate (Whitman and Greer, 2007b; Batista-Brito et al., 2008).

**Figure 5.**
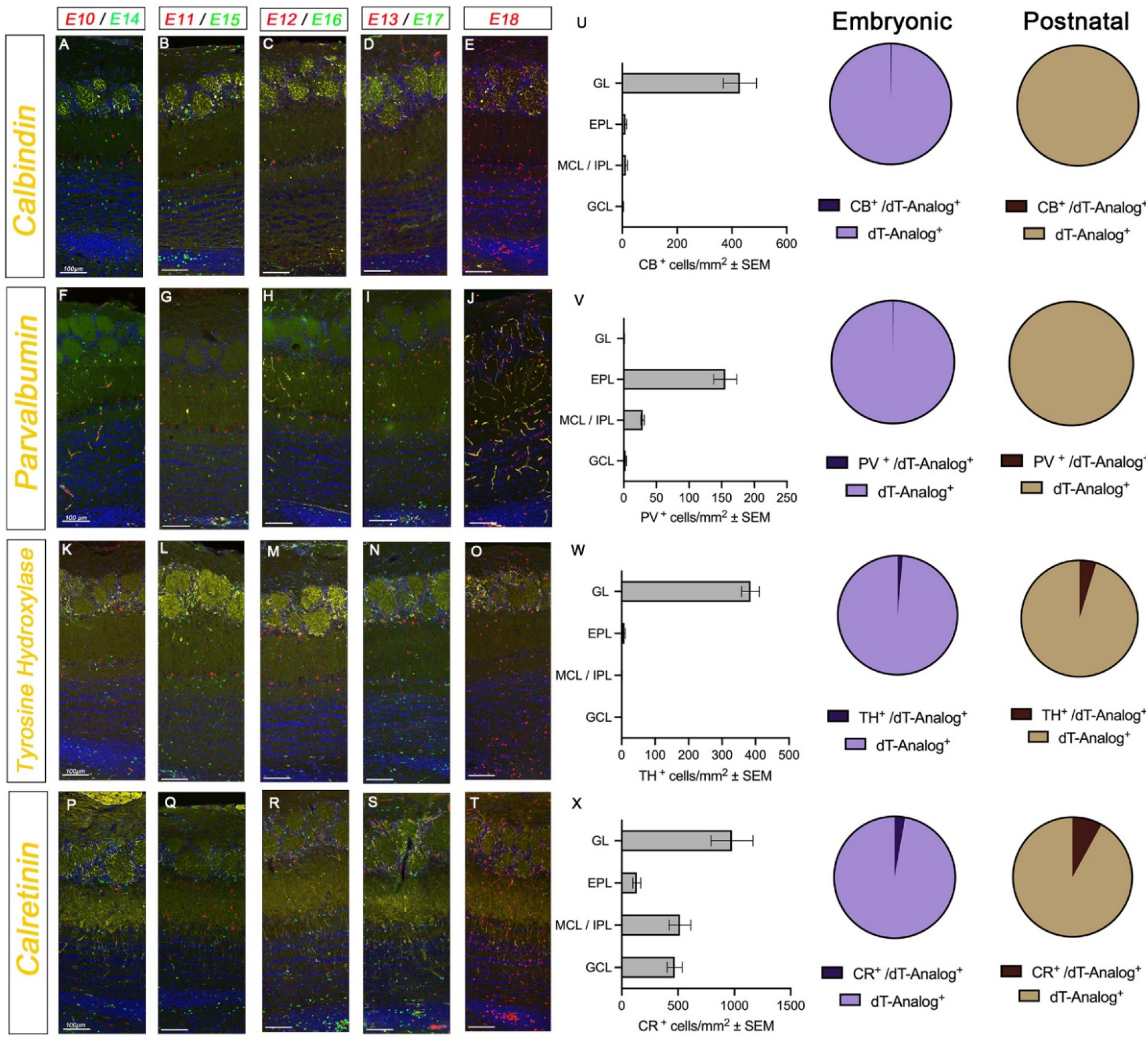
Characterization of EGINs and PGINs phenotypes. Phenotypes are identified using the common interneuron markers: CB, PV, TH, CR. Panels are representative images of OB stained with IdU^+^ cells (red), CldU^+^ cells (green) and INs markers in yellow. Nuclei counterstained with DAPI (blue). (**A-E)** Examples of immunostaining for CB across the OB layers. (**F-J)** Examples of immunostaining for PV across the OB layers. (**K-O)** Examples of immunostaining for TH across the OB layers. (P-T) Examples of immunostaining for CR across the OB layers. (**U-X)** Quantification of the total number of CB, PV, TH and CR cells by layer (left) and representation of colocalization with the thymidine analogs as pie charts from EGINs (purple) and PGINs (brown) (right). We do not observe a percentage higher than 8% (CR in PGINs) for any of the quantified cells. Abbreviations: CB: calbindin; CR: calretinin; dT-Analog: thymidine analog; EPL: external plexiform layer; GCL: granule cell layer; GL: glomerular layer; MCL/IPL: mitral cell layer/internal plexiform layer; PV: parvalbumin; TH: tyrosine hydroxylase. Scale bars: 100 µm.

### Migration of EGINs during embryonic stages

Finally, we sought to investigate the migration and differentiation of EGINs during embryonic stages of development. To achieve this, we use IUE to target EGIN progenitors in the LGE where OB INs are embryonically generated (Toresson and Campbell, 2001; Wichterle et al., 2001; Guo et al., 2019). Progenitor cells were permanently labeled using a multicolor technique based on the piggyBac transposon at E11 (Fig. 6A) (Martin-Lopez et al., 2019a; Martin-Lopez et al., 2019b).

**Figure 6.**
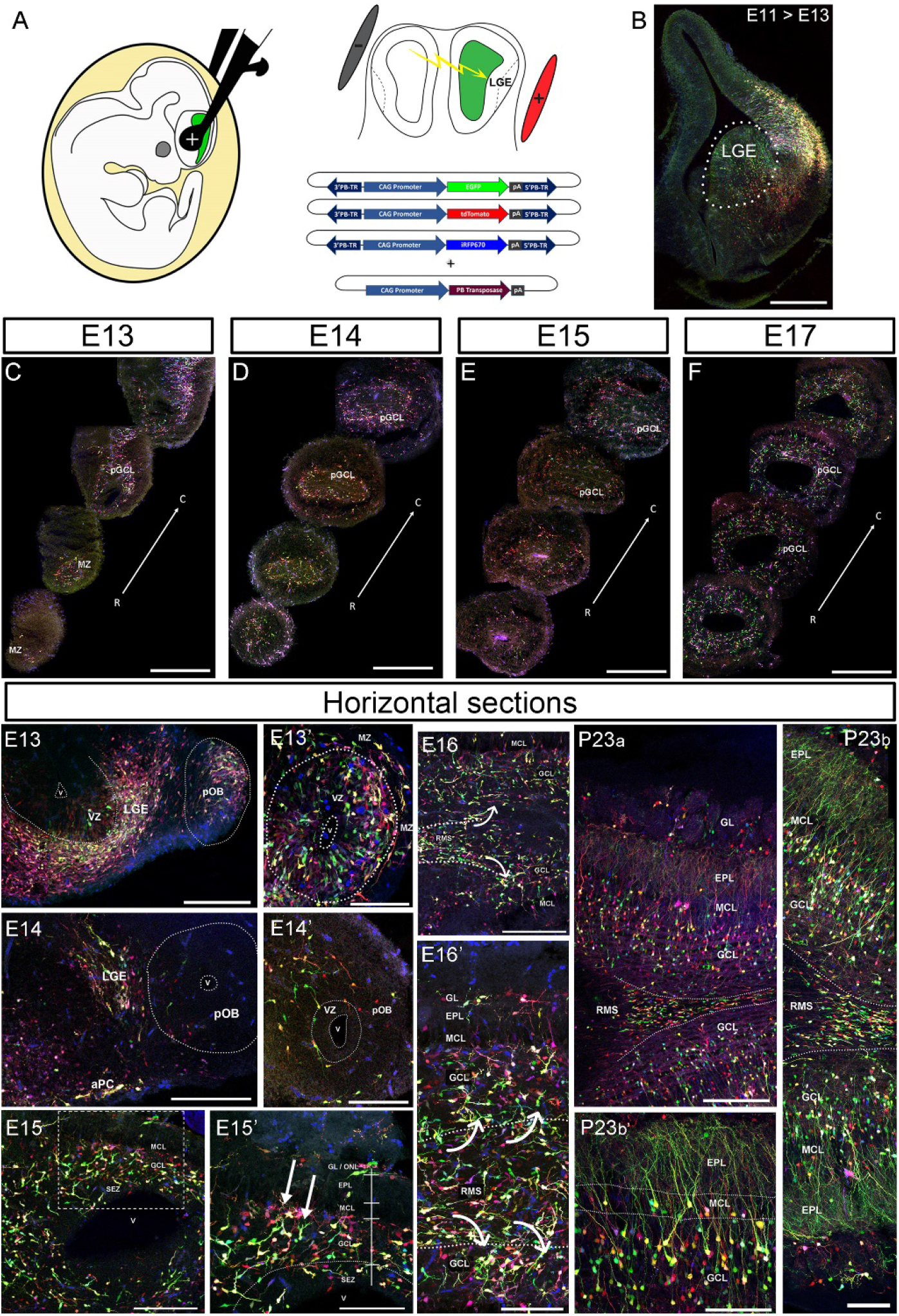
Migration and differentiation of EGINs at embryonic stages. **(A)** Diagram representing the electroporation area targeting the LGE and the piggyBac plasmids used for the procedure. **(B)** Representative image of a coronal brain section of an E13 embryo showing the labeling of EGINs progenitors in the LGE. **(C-F)** Representative images of the primordial OBs from electroporated mice across the rostral to caudal axis. Prospective GCL is seen occupied by neuroblasts at all ages. **(Horizontal Sections)** Representative images of embryonic sections showing the migration and differentiation of EGINs. Progenitor cells are arriving to the pOB as early as E13 filling the MZ **(E13, E13’)**. At E14 EGIN progenitors extend leading process while migrating to the pOB **(E14, E14’)**. At E15 we observe the first evidence of apical dendrites differentiation of EGINs in the GCL and GL of the pOB **(E15, E15’ arrows)**. At E16 the apical dendrites from EGINs are seen to extend from the GCL to the EPL, and neuroblasts are seen clustered forming a chain of migrating neuroblasts that form the first evidence of a RMS **(E16-E16’, curved arrows)**. At adult stages (P23), the morphology of the fully formed OB is evident, showing EGINs in the different layers exhibiting complex dendritic arbors that extend towards the EPL. RMS is clearly distinctive at this age (P23, P23a, P23b). Abbreviations: aPC: anterior piriform cortex; C: caudal; EPL: external plexiform layer; GCL: granule cell layer; GL: glomerular layer; LGE: lateral ganglionic eminence; MCL: mitral cell layer; MZ: marginal zone; ONL: olfactory nerve layer; pGCL: prospective granule cell layer; pOB: prospective olfactory bulb; R: rostral; RMS: rostral migratory stream; SEZ: subependymal zone; V: ventricle; VZ: ventricular zone. Scale bars: B: 200μm; E13, E14, E14’, E15, E16, P23a: 100μm; E13’, E15’, E16’, P23b, P23b’: 50μm.

Our results in coronal sections showed that precursors of EGINs were already present in the marginal zone (MZ) of the primordium of the OB as early as E13, even in the most rostral sections (Fig. 6C). As embryonic development progressed, labeled cells were seen to increase in both numbers and complexity, predominantly accumulating in the developing nascent GCL (Fig. 6D-F). To study the migration and differentiation of EGIN precursors, we used a different set of embryos that were sectioned horizontally (Fig. 6, Horizontal). Here, we observed that EGIN precursors were located predominantly in the ventricular zone (VZ) of the LGE, which was adjacent to the MZ of the prospective OB (Fig. 6 E13, E13’). These cells surrounded the VZ and exhibited a migratory shape of undifferentiated neuroblasts, extending a forward process known as the leading process (Fig. 6 E13, E13’). Similar behavior was observed at E14, where progenitor cells were still randomly distributed next to the LGE (Fig. 6 E14, E14’). One day later at E15, we observed the first evidence of EGIN differentiation, where the leading processes were transformed into a dendritic tuft that turned toward the surface of the OB (Fig. 6 E15, E15’ arrows). Some EGIN precursors began to migrate into the primordial GL and neuroblasts were visible clustering next to the OB. At E16 the dendritic tufts from differentiating EGINs appeared visibly more mature and begin to resemble an apical dendrite (Fig. 6 E16’ in EPL). For the first time at this age, we noticed a chain of clustered migrating neuroblasts as the ventricle retracts from the OB, forming a visible RMS (Fig. 6 E16, E16’ arrows). Finally, at P23 we observed a completely formed OB, where EGINs were distinctively differentiated across the OB layers, extending apical dendrites that arborized in the EPL to contact M/Tc (Fig. 6 P23a and b, b’). Most EGINs accumulated in the GCL, surrounding the neuroblast chain that now fully formed the RMS (Fig. 6 P23a and b). Importantly, our results showed that using the piggyBac transposon was an improved method to detect the formation of the RMS embryonically, and allowed us to show neuroblasts at a much higher resolution level compared to those using only immunohistochemistry (Pencea and Luskin, 2003).

## Discussion

The OB constitutes an extraordinary plastic environment characterized by a constant supply of neurons responsible for important adaptative functions (Sailor et al., 2017). These cells are INs that are generated in a process that begins early during the embryonic development and persists until the end of the life of an individual (Treloar et al., 2010; Tufo et al., 2022). Embryonically, OB neurogenesis takes place in the LGE, which slowly disappears and turns into a small neurogenic zone surrounding the walls of the lateral ventricles. Referred to as the subventricular zone (SVZ) postnatally, it remains as a proliferative region of the brain. From here, OB neuroblasts travel towards the OB in a chain of migrating neuroblasts named the rostral migratory stream (RMS) (Wichterle et al., 2001; Whitman and Greer, 2009; Lim and Alvarez-Buylla, 2016; Lledo and Valley, 2016; Tufo et al., 2022). Once these neuroblasts arrive and differentiate into OB INs, their final fate is directly influenced by their generation date and environment (Lemasson et al., 2005; Batista-Brito et al., 2008; Lledo et al., 2008; Galliano et al., 2018; Kim et al., 2020; Liberia et al., 2024). The importance of studying the relationships between the OB interneuron phenotypes with their generation dates is critical to understanding the morphology and function of the OB circuitry. Here, we focused our work only on those OB INs that are produced at embryonic stages, that we have broadly named “Embryonic Generated Interneurons” or EGINs. Our goal is to provide insight into the EGIN developmental dynamics and their final phenotypes once they are already incorporated into the OB circuits. The birthdates of EGINs were studied by injections of thymidine analogs in pregnant mouse females, and their final fate characterized by immunohistochemistry. We were also interested in analyzing neurogenic gradients along the lateral-to-medial axis, since this particular gradient influences the development of neurons in the tubular striatum (a.k.a. olfactory tubercle) and the OB projection neurons M/Tc (Imamura et al., 2011; Imamura and Greer, 2015; Martin-Lopez et al., 2019b). Finally, we carried out experiments using IUE to label EGIN progenitors in the LGE to follow their trajectory and differentiation in the developing OBs.

The presence of INs generated embryonically in the adult OB has been studied previously, but their final destinations, differentiation and migratory routes during the embryogenesis have not been fully characterized. For instance, pioneering work in mice using injections of tritiated thymidine show, as we do, that there is a spike in EGINs generation at E18 (Hinds, 1968b). However, Hinds did not find any neurogenic gradient from granule cells as we did; as well he reported a sharp increase in granule cells from E15 onwards, that we could not replicate. Although we do not have an explanation for these discrepancies, we can speculate with differences in the methodological sensitiveness between tritiated thymidine and immunohistochemical detection of thymidine analogs. We do agree with Hinds nonetheless that the day of birth for OB INs may be arbitrary due to the extremely long generation period of these cells (Hinds, 1968b). This may suggest that EGINs generated at E18 are more closely related to postnatal INs (PGINs) than the embryonic ones. Further investigation is needed to confirm this possibility.

It was striking to find an EGINs lateral-to-medial neurogenic gradient as occurs in other areas of the olfactory system such as the tubular striatum or the anterior olfactory nucleus (Creps, 1974; Martin-Lopez et al., 2019b). This type of gradient has been reported before in other pallial brain structures (Pierani and Wassef, 2009), but this is the first time that it is demonstrated in populations of INs. This is particularly revealing since mitral cells, which are the main projection neurons of the OB, follow a developmental gradient that is reversed from that observed with the EGINs: mitral cells populating medial OB regions are produced prior to the ones from the lateral regions (Imamura and Greer, 2015).

Understanding how two closely related neurons that collectively establish the OB circuitry can follow inverted generation gradients during development poses a provocative challenge. However, it is reasonable to speculate with the hypothesis that EGINs follow a pre-established genetic program that is shared by other pallial neurons, considering that all EGINs come from the dorsal region of the LGE that has a pallial origin (Li et al., 2018; Kuerbitz et al., 2021).

In the context of EGINs differentiation, we found fewer EGINs differentiated into the commonly identifiable IN subpopulations, positive for the CB, PV, TH, CR markers, compared to a prior report (Batista-Brito et al., 2008). One potential explanation is that our numbers could be undercounted due to a different and more extensive subpopulation of EGINs than those described by Batista-Brito and colleagues (Batista-Brito et al., 2008). Thus, these authors used a Dlx1/Dlx2 mouse model to track embryonic INs in the adult OB. The Dlx1/Dlx2 mouse model identifies only those OB INs that are part of the dorsal domain of the LGE (dLGE) that expressed these transcription factors (the Dlx1^+^/Dlx2^+^ domain) (Bulfone et al., 1998; Long et al., 2007). However, it excludes IN sources outside of the dLGE such as the Emx1^+^ and Dlx5^+^/Dlx6^+^ regions of the pallium and septum (Kohwi et al., 2007; Qin et al., 2017), which may also give rise to OB INs. Because we used injections of thymidine analogs that did not distinguish between different EGINs generative zones, the numbers in our analyses may reflect a more diverse population of EIGNs that then tend to be more diverse than in prior studies. In addition, at the time of our analysis at P21, the differentiation of EGINs may be still incomplete. Neurons such as those expressing TH, that are generated embryonically at E12, have been found to almost quadruple their differentiation rate from the 1^st^ to the 6^th^ month of postnatal survival (Galliano et al., 2018).

In summary, we show here that the production, migration, differentiation, and maturation of EGINs seems to be a very dynamic process that is affected by both the environment and the age of the animals-highlighting the complexity of the circuitry in the OB. We found that embryonically generated interneurons of the olfactory bulb (EGINs) are found across all layers of the OB exhibiting an average diameter of 7 μm. In addition, EGINs display a lateral-to-medial neurogenic gradient that concentrates in the EPL at E11 and in the MCL/IPL at E12-E13. Within the GCL, EGINs show a superficial-to-deep laminar distribution at most gestational stages excepting E16 and E18. Unexpectedly, a small percentage of EGINs expressed any of the most common INs markers for CB, PV, TH and CR, suggesting that they are still unknown subpopulations of OB interneurons. Finally, we found that EGINs migrate from the LGE towards the prospective OB as early as E13, extending differentiated apical dendrites at E15, and organizing in a chain of migrating neuroblasts constituting the embryonic RMS at E16. At adult stages, most EGINs are morphologically differentiated into OB INs such as granule and periglomerular neurons.

## Acknowledgements

Support provided by NIH-NIDCD 013791, NIH-NIDCD 016851 and NIH-NIDCD 017989 to CAG. The authors express their appreciation to Christine Kaliszewski for technical assistance.

## Conflict of interest statement

All authors declare no competing financial interests.

## Notes

### Competing Interest Statement

The authors have declared no competing interest.

